# *Staphylococcus aureus* inhibits the NLRP3 inflammasome in macrophages during the early phases of intracellular infection, but not the late phases

**DOI:** 10.1101/2025.09.24.678325

**Authors:** Saumya Bhagat, Khushpreet Kaur, Luke O’Connor, Chun Wang, Yongjia Li, Gaurav Swarnkar, Kunjan Khanna, James E. Cassat, Deborah J. Veis, Gabriel Mbalaviele

## Abstract

*Staphylococcus aureus* (*S. aureus*) is a highly virulent pathogen responsible for chronic infections such as osteomyelitis. Although its interaction with the host immune system has been widely studied, the specific role of inflammasomes in regulating the infection within macrophages remains unclear. We investigated this question using bone marrow-derived macrophages infected with *S. aureus* and observed a significant reduction in intracellular bacterial load beginning at 18 hours post-infection (hpi), which continued through 96 hpi. Notably, robust activation of the NLRP3 inflammasome—including inflammasome assembly, IL-1β and GSDMD maturation, and pyroptosis—occurred only after 18 hpi. This led us to hypothesize that *S. aureus* suppresses inflammasome activation during early infection. Supporting this, infected BMDMs failed to respond robustly to LPS and nigericin up to 18 hpi, with partial recovery at later timepoints, suggesting that *S. aureus* initially inhibits NLRP3 signaling to persist within macrophages but is later counteracted by the host response.

## INTRODUCTION

*Staphylococcus aureus (S. aureus)* is a highly versatile Gram-positive pathogen responsible for a wide range of infections, including osteomyelitis^1^. Commonly considered as an extracellular pathogen, *S. aureus* also survives within host cells^2^. Its ability to infect, survive, and propagate in bone microenvironment within various cell types including macrophages, osteoclasts, neutrophils, and mesenchymal cells is central to the pathogenicity of chronic osteomyelitis^3^. The ability of *S. aureus* to thrive as a pathogen can be largely attributed to its extensive array of virulence factors, which allow the bacterium to efficiently extract nutrients from its host while evading both innate and adaptive immune defenses^4,5^. Macrophages are pivotal in the immune responses against pathogens, serving as the first line of defense through phagocytosis and the initiation of inflammatory responses. They are equipped with a variety of pattern recognition receptors such as Toll-like receptors and NOD-like receptors (NLRs), which are essential for detecting pathogen-associated molecular patterns (PAMPs)^6^. They are also highly efficient at phagocytosing *S. aureus* which upon internalization, are enclosed in phagosomes, which fuse with lysosomes to form phagolysosomes^7^. The acidic environment within the phagolysosome, along with reactive oxygen species (ROS) and reactive nitrogen species, contributes to the destruction of the bacteria^8,9^. Additionally, macrophages produce a wide range of cytokines and chemokines in response to *S. aureus* infection such as TNF-α, IL-1β, and IL-6, which help recruit additional immune cells to the site of infection and amplify the inflammatory responses^10,11^. However, *S. aureus* has evolved several strategies to evade killing within macrophages. For example, it produces catalase, which neutralizes ROS, and proteins that inhibit phagosome-lysosome fusion, allowing the bacterium to survive and replicate intracellularly^12^. Given the immune functions of macrophages, it stands to reason that evasion of macrophage-dependent killing is required to successfully establish and maintain a productive infection.

The inflammasomes assembled by NLR family pyrin domain containing 3 (NLRP3) and absent in melanoma 2 (AIM2) play critical roles in host defense against *S. aureus* infection. NLRP3 detects bacterial toxins and lipoproteins which cause membrane damage, lysosomal rupture, and ROS production, thereby triggering inflammasome activation to control the infection^13–15^. The AIM2 inflammasome is particularly important in fighting intracellular bacterial pathogens such as *S. aureus, L. monocytogenes, F. tularensis* and *M. tuberculosis* upon detection of DNA in the cytoplasm, which originates from lysed bacteria^16,17^. Although activated through distinct mechanisms, both NLRP3 and AIM2 assemble macromolecular protein complexes that include apoptosis-associated speck-like protein containing a CARD (ASC) and caspase-1^18–20^. Caspase-1 not only processes pro-IL-1β and pro-IL-18 into their active forms, but also cleaves gasdermin D (GSDMD), generating amino-terminal fragments that form plasma membranes pores through which these cytokines are secreted^21–23^.

Despite extensive research, several key elements regarding *S. aureus* and inflammasome activation remain unclear, including the precise mechanisms by which *S. aureus* modulates inflammasome activity during acute and chronic infections, the contribution of alternative inflammasome pathways beyond NLRP3 in the detection and response against *S. aureus*, and the balance between protective immune responses and pathological inflammation in diseases such as osteomyelitis and sepsis. Filling this knowledge gap is essential for designing targeted therapies that enhance immunity while minimizing tissue damage.

In this study, we studied the interactions between *S. aureus* and BMDMs. We identified NLRP3 but not AIM2 as the primary sensor of this bacterium in BMDMs. We also found that *S. aureus* initially suppresses NLRP3 inflammasome activation, but as the infection progresses, BMDMs overcome this inhibition, at least partially.

## RESULTS

### The killing of *S. aureus* by BMDMs is time- and NLRP3 inflammasome-dependent

FBS supports the survival, growth, and immune responses of phagocytes, including macrophages^24,25^. To evaluate the impact of serum supplementation on the ability of BMDMs to clear *S. aureus* infection, WT BMDMs were cultured in media containing 0, 1, or 10% FBS and subsequently infected with Ti3 *S. aureus* strain (hereafter referred to as *S. aureus*) at a multiplicity of infection (MOI) of 100. The CFU assay showed a decrease in bacterial load at 18 hpi compared to 1.5 hpi across all conditions, with significantly greater clearance observed with 10% FBS supplementation than with 0% FBS, underscoring the impact of serum on bacterial elimination (Fig 1a).

**Fig 1:**
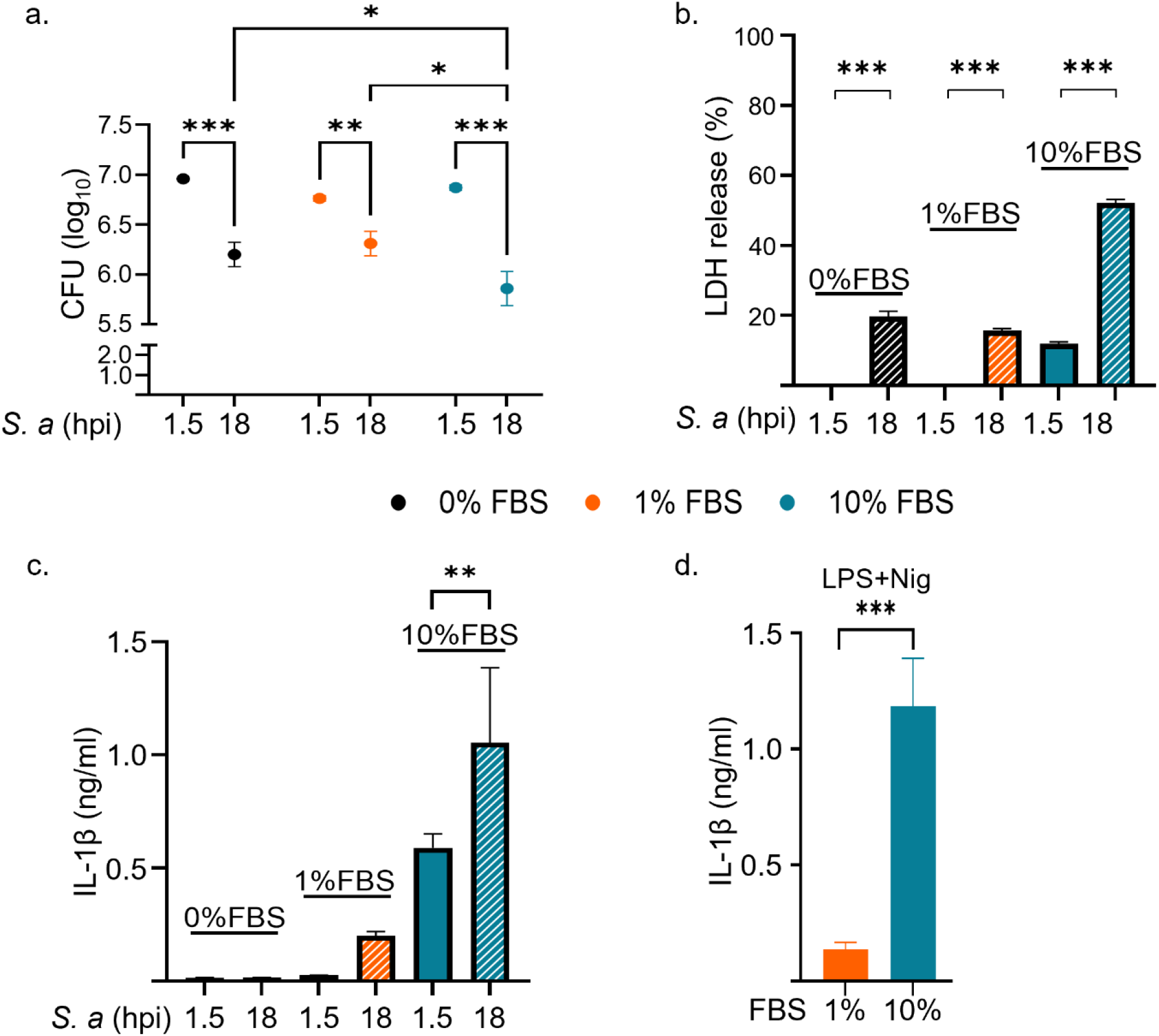
*S. aureus* is more efficiently killed by BMDMs cultured in media containing a high concentration of FBS. WT BMDMs were infected with *S. aureus* (*S. a*) under varying serum conditions, 0% FBS (serum starved), 1% FBS (reduced serum) and 10% FBS (normal serum) and analyzed post-infection. a. CFUs from lysates of infected BMDMs were quantified at 1.5 hpi, immediately after gentamicin exposure, or after an additional 16.5 h in culture media (18 hpi). b. LDH release in the conditioned media was measured using the LDH kit. c. IL-1β secretion in the conditioned media were evaluated by ELISA. d. Uninfected BMDMs treated with 1% or 10% FBS were primed with LPS for 3 h followed by 45 min of nigericin treatment. Data are means ± SD from experimental triplicates and are representative of at least two independent experiments. *p<0.05; **p<0.01; ***p<0.001 by two-way ANOVA (a), one-way ANOVA (b, c) with Bonferroni’s *post hoc* test or Student’s *t* test (d).

The inflammasomes are induced by *S. aureus* and are responsible for the maturation and secretion of IL-1β, and they can also induce pyroptosis^13,26^. Therefore, we analyzed the secretion of IL-1β and lactate hydrogenase (LDH; a readout of pyroptosis). LDH release was significantly higher at 18 hpi compared to 1.5 hpi across all conditions, with the maximal response observed in cultures containing 10% FBS (Fig 1b). By contrast, IL-1β levels were only detected in cells exposed to 1% or 10% FBS, with the highest response observed in the 10% FBS group (Fig 1c). Notably, the levels of IL-1β triggered by *S. aureus* were comparable to those induced by LPS and nigericin, which served as positive controls (Fig 1d).

Given that BMDMs exhibit optimal control of *S. aureus* infection, LDH release, and IL-1β secretion in the presence of 10% FBS, we leveraged this condition to assess their ability to clear this bacterium and secrete IL-1β over time, up to 96 hpi. The number of CFUs declined in a time-dependent manner (Fig 2a), a pattern that inversely correlated with the secretion of IL-1β, which peaked at 24 hpi (Fig 2b), and LDH release (Fig 2c). These data suggest that inflammasomes may be involved in the ability of BMDMs to clear *S. aureus* infection, a response that is robust at later times post-infection.

**Fig 2:**
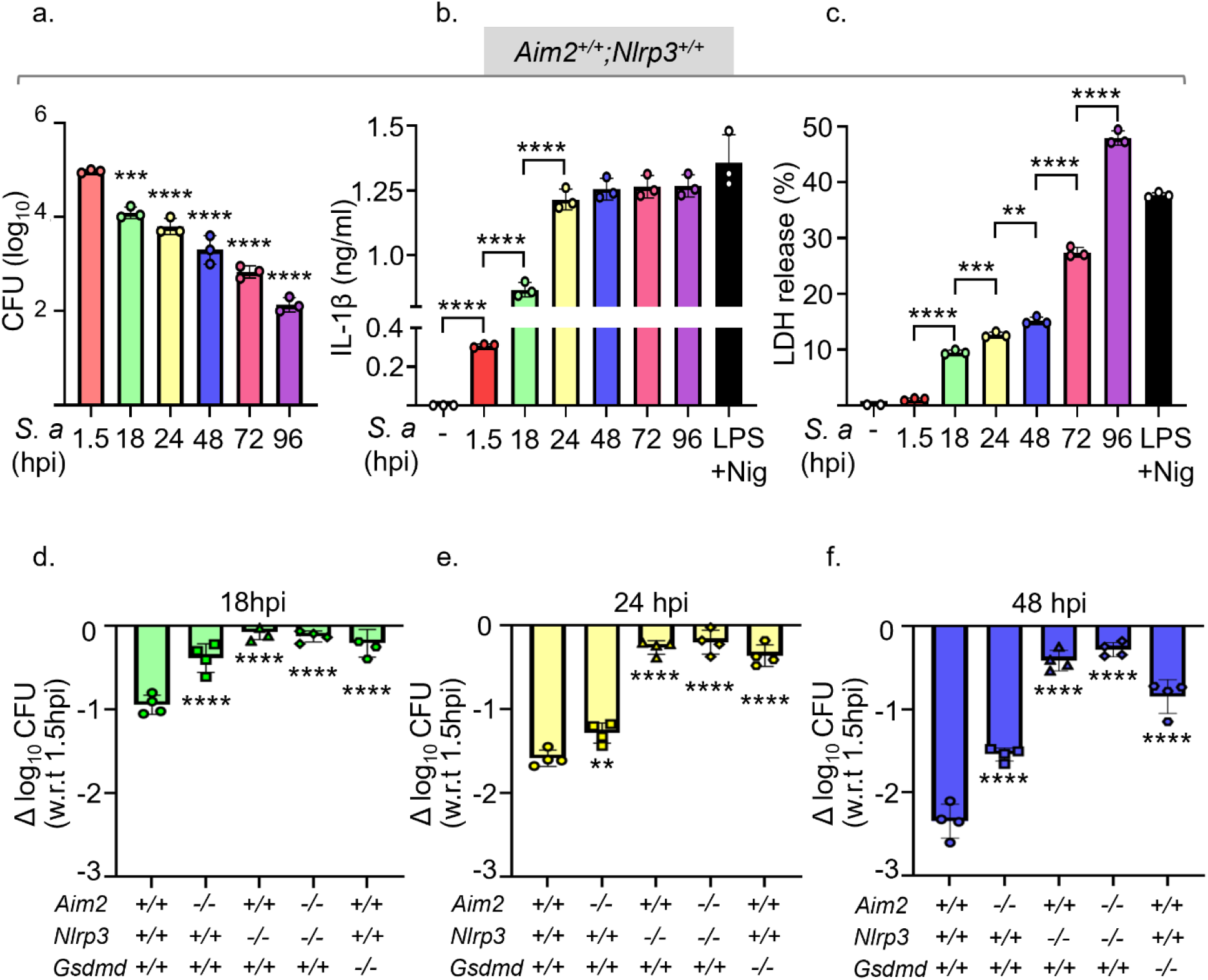
The killing of *S. aureus* by BMDMs is time- and NLRP3 inflammasome-dependent. *Aim2*^*+/+*^*;Nlrp3*^*+/+*^, *Aim2*^*+/+*^*;Nlrp3*^*-/-*^, *Aim2*^*-/-*^*;Nlrp3*^*+/+*^, *Aim2*^*-/-*^*;Nlrp3*^*-/-*^, and *Gsdmd*^*-/-*^ BMDMs were infected with *S. aureus* for up to 96 h. Cells treated sequentially with LPS for 3 h and nigericin for 45 min served as positive controls. Timepoint analysis of WT BMDMs was assessed by a. CFU, b. IL-1β secretion. c. LDH release. d-f. CFU assessment was also done for knockout genotypes and compared to WT and data were normalized to 1.5 hpi internalization timepoint. n=3 biological replicates. Data are means ± SD from experimental triplicates. *p<0.05; **p<0.01; ***p<0.001; ****p<0.0001 by one-way ANOVA with Bonferroni’s *post hoc* test.

To investigate the role that inflammasomes play in the fate of *S. aureus* in our experimental models, we scored CFUs in BMDMs expressing or lacking AIM2 or NLRP3, which are both well known for sensing this bacterium^27–29^. The ability of *Aim2*^*+/+*^*;Nlrp3*^*+/+*^ BMDMs to clear *S. aureus* infection increased in a time-dependent manner (Fig 2d-f), consistent with the results shown in Fig 2a. Although the killing of *S. aureus* by *Aim2*^*-/-*^*;Nlrp3*^*+/+*^ BMDMs was slight impaired at 18 hpi, this response was almost fully restored in these cells at later timepoints (Fig 2d-f; Supplementary Fig 1a). In contrast, *Aim2*^*+/+*^*;Nlrp3*^*-/-*^ BMDMs and *Aim2*^*-/-*^*;Nlrp3*^*-/-*^ counterparts were unable to eliminate this bacterium (Fig 2d-f; Supplementary Fig 1a). Comparable results were obtained when the fate of *S. aureus* in WT and mutant BMDMs was measured by flow cytometry (Supplementary Fig 1b-c). GSDMD is processed alongside pro-IL-1β upon AIM2 or NLRP3 inflammasome activation^22^. Therefore, we also determined the impact of GSDMD deficiency on the fate of *S. aureus* in BMDMs. The number of CFUs did not change significantly over time in *Gsdmd*^*-/-*^ BMDMs (Fig 2d-f; Supplementary Fig 1a). These results suggest that the NLRP3 inflammasome-GSDMD axis plays the key role in the elimination of *S. aureus* by BMDMs.

We also monitored the kinetics of IL-1β secretion and LDH release during the progression of BMDM infection by *S. aureus*. The levels of secreted IL-1β by WT BMDMs were low at 1.5 hpi, but increased over time, reaching a maximum at 48 hpi (Fig 3a-f). Consistent with CFU results, while the levels of IL-1β were only marginally altered in *Aim2*^*-/-*^*;Nlrp3*^*+/+*^ BMDMs compared to *Aim2*^*+/+*^*;Nlrp3*^*+/+*^ cells, they were markedly reduced at all timepoints in *Nlrp3*^*-/-*^, *Aim2*^*-/-*^*;Nlrp3*^*-/-*^, or *Gsdmd*^*-/-*^ BMDMs (Fig 3a-f). Likewise, the release of LDH by *Aim2*^*+/+*^*;Nlrp3*^*+/+*^ and *Aim2*^*-/-*^*;Nlrp3*^*+/+*^ BMDMs, which was minimal at 1.5 hpi, increased progressively thereafter, and then plateaued at 48 hpi (Fig 3g-l). These outcomes were nearly suppressed in *Aim2*^*+/+*^*;Nlrp3*^*-/-*^, *Aim2*^*-/-*^*;Nlrp3*^*-/-*^, or *Gsdmd*^*-/-*^ BMDMs (Fig 3g-l). Collectively, these findings suggest that the NLRP3 inflammasome is the primary sensor of *S. aureus*, with its bactericidal activity increasing as the infection progresses.

**Fig 3:**
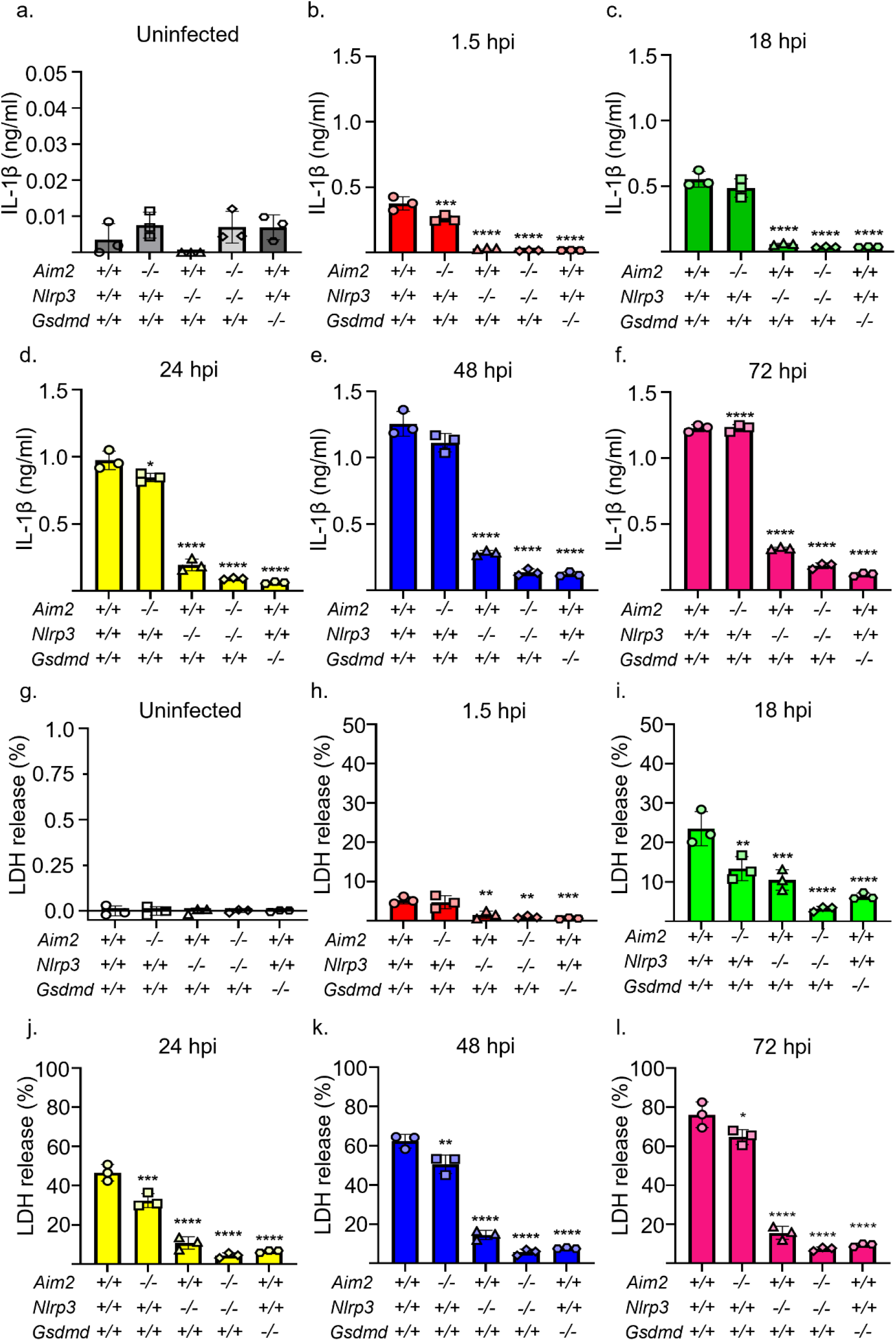
*S. aureus*-induced IL-1β secretion and LDH release are time- and NLRP3 inflammasome-dependent. *Aim2*^*+/+*^*;Nlrp3*^*+/+*^, *Aim2*^*+/+*^*;Nlrp3*^*-/-*^, *Aim2*^*-/-*^*;Nlrp3*^*+/+*^, *Aim2*^*-/-*^*;Nlrp3*^*-/-*^, and *Gsdmd*^*-/-*^ BMDMs were left uninfected or infected with *S. aureus* for up to 96 h. The supernatants were analyzed for IL-1β secretion and LDH release. Uninfected cells sequentially treated with LPS for 3 h and nigericin for 45 min were used as positive controls. a-f. IL-1β secretion. g-l. LDH release. *n* = 3 technical replicates, representative of 3 biological replicates. Data are expressed as means ± SD. **p<0.01; ***p<0.001; ****p<0.0001 by one-way ANOVA with Bonferroni’s *post hoc* test.

### *S. aureus* suppresses the NLRP3 inflammasome in BMDMs during the early, but not late phases of infection

As mentioned above, upon activation, NLRP3 undergoes oligomerization and recruits ASC, which then forms polymers. These macrostructures, known as ASC specks, can be visualized as foci under a fluorescence microscope^18^. Uninfected BMDMs from *Asc-citrine* mice readily formed ASC specks in the presence of LPS and nigericin (Fig. 4a). By contrast, ASC specks in *S. aureus*-infected BMDMs were detected only after 18 hpi and were present in approximately 5% of cells (Fig. 4b, c). Given the delay in ASC speck formation, we hypothesized that the NLRP3 inflammasome was suppressed during the initial stages of infection. To evaluate this hypothesis, we examined the activation of this inflammasome in response to LPS and nigericin in both uninfected and infected cells. While approximately 60% of uninfected BMDMs treated with LPS and nigericin formed ASC specks, this response was blunted in infected cells up to 18 hpi and was only partially restored at 24, 48, and 72 hpi (Fig. 4a, b, c). Similar trends of inflammation activation were observed when we monitored the activated state of caspase-1 (Supplementary Fig 2a, b).

**Fig 4:**
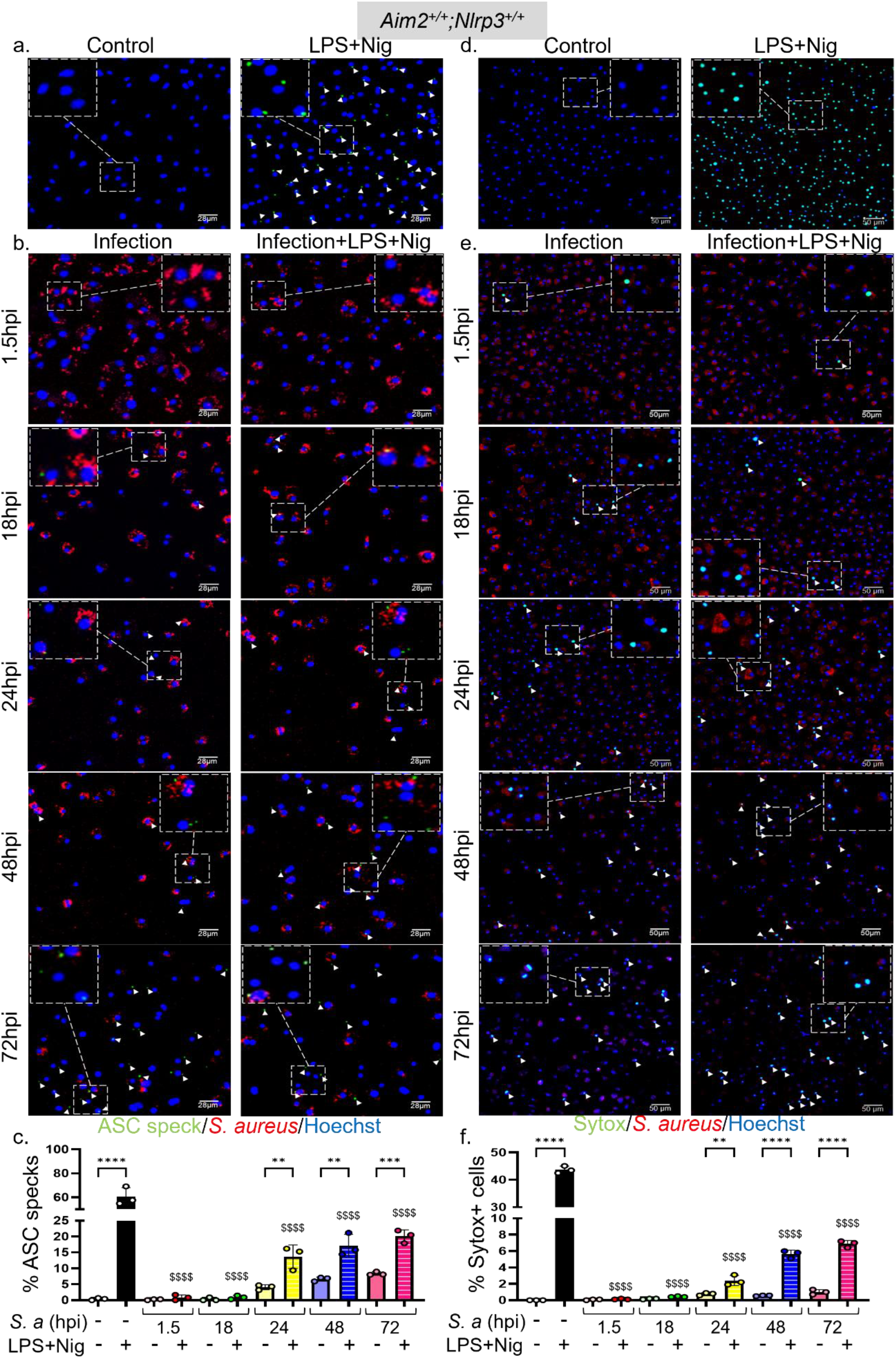
*S. aureus* suppresses ASC speck formation and Sytox Green uptake during the early phases of infection of BMDMs, but to a lesser extent during the late phases. BMDMs from *ASC-citrine* mice (to assess ASC speck formation) and WT mice (to measure Sytox Green uptake) were infected with *S. aureus* for up to 72 h. Uninfected left untreated or treated sequentially with LPS for 3 h and nigericin for 45 min of nigericin were used as controls. The cells were incubated with Hoechst 33342 (1 µg/mL) for 15 min after LPS treatment. a, b. ASC specks were visualized under fluorescence microscopy and c. quantified using ImageJ. d, e. Sytox Green was added to the cultures 10 min prior to imaging by confocal microscopy and f. the uptake was quantified using ImageJ. Data are means ± SD from experimental triplicates and represent at least two independent experiments. **p<0.01; ***p<0.001; ****p<0.0001 by one-way ANOVA with Bonferroni’s *post hoc* test. Comparison of LPS+Nig positive control with infection samples treated with LPS+Nig shown as $$$$ p<0.0001 by one-way ANOVA.

The NLRP3 inflammasome promotes the uptake of the DNA-binding dye Sytox Green^30^. Accordingly, the kinetics of Sytox Green uptake and, to a lesser extent, IL-1β secretion induced by LPS and nigericin closely mirrored the formation of ASC specks (Fig. 4d, e, f; Fig 5a). Immunoblotting analysis of cell lysates indicated that the expression of NLRP3 and pro-IL-1β, but not of GSDMD, was maximally enhanced by *S. aureus* at 24 hpi before declining at later timepoints (Fig 5b). Cleaved GSDMD amino-terminal fragments (p30) and mature IL-1β (p17) were readily detected in samples from LPS and nigericin-treated uninfected and infected BMDMs (Fig 5b). Unexpectedly, *S. aureus* did not promote GSDMD cleavage and appeared to induce the generation of IL-1β fragments of various molecular weights (Fig 5b). Technical issues, including cell loss and overestimation of protein concentrations, could account for the lack of protein detection at the 72 hpi timepoint. In any case, these findings suggest that *S. aureus* initially suppresses NLRP3 inflammasome activation, delaying ASC speck formation and downstream responses, which are partially restored at later stages of infection.

**Fig 5:**
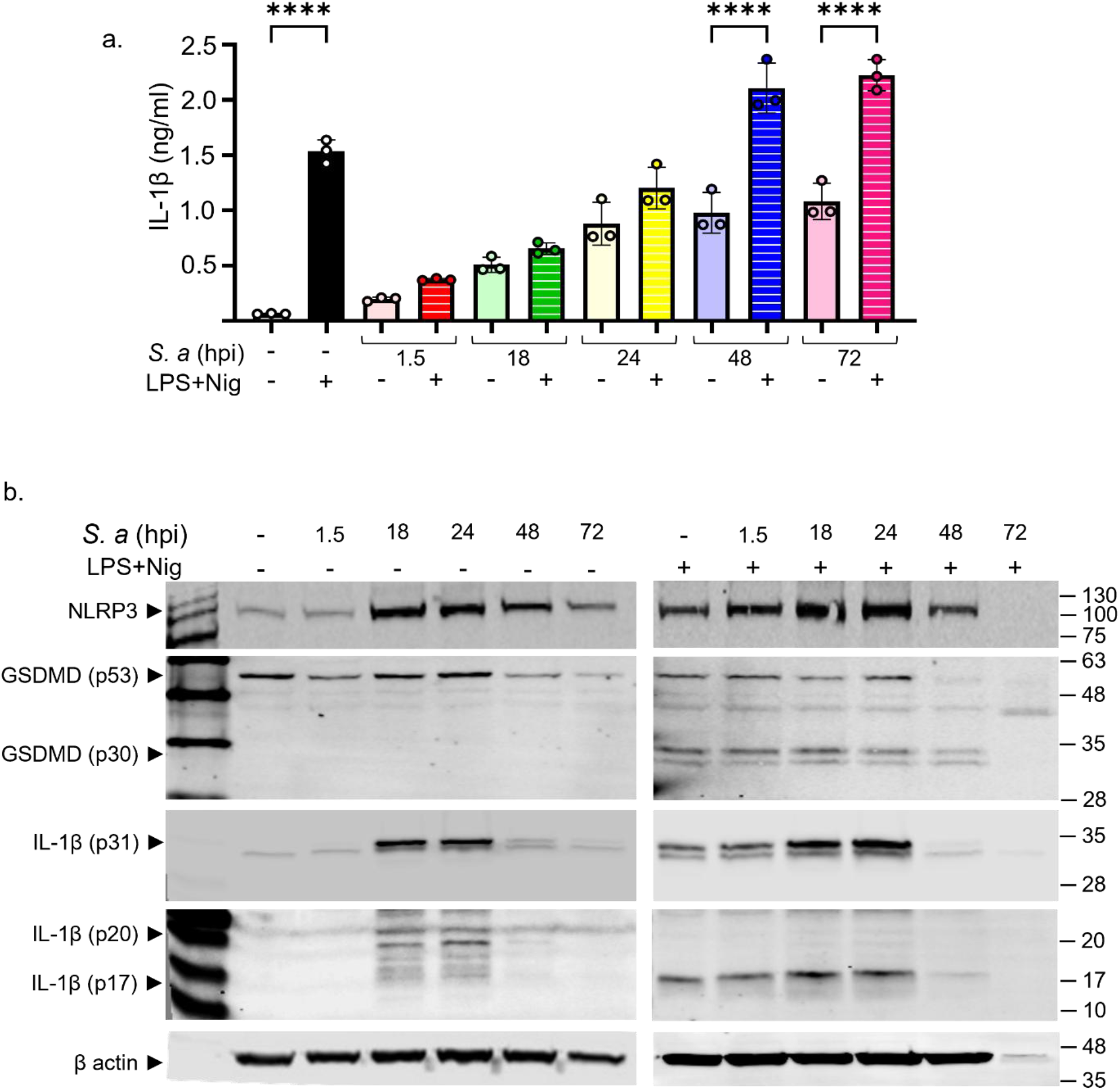
*S. aureus* affects IL-1β and GSDMD maturation. Uninfected and WT BMDMs infected with *S. aureus* were left untreated or primed with LPS for 3 h, then treated with nigericin for 45 min. a. IL-1β. b. Immunoblotting analysis of cell lysates. Data are means ± SD from experimental triplicates and represent at least two independent experiments. ****p<0.0001 by one-way ANOVA with Bonferroni’s *post hoc* test.

To further demonstrate that *S. aureus* primarily modulates the inflammasome assembled by NLRP3, but not AIM2, we examined the activation of these inflammasomes in response to LPS and nigericin in both uninfected and infected BMDMs, assessing each sensor individually and in combination. We also analyzed the impact of GSDMD deficiency on *S. aureus*-induced outcomes. We found that the formation of ASC specks induced by LPS and nigericin was similarly modulated *S. aureus* in the presence of NLRP3 but completely abolished in its absence, regardless of AIM2 or GSDMD (Fig 6a; Supplementary Fig 3). Accordingly, the uptake of Sytox Green, although marginal, was observed only in *Aim2*^*-/-*^*;Nlrp3*^*+/+*^ BMDMs, but not in *Aim2*^*+/+*^*;Nlrp3*^*-/-*^, *Aim2*^*-/-*^*;Nlrp3*^*-/*^, or *Gsdmd*^*-/-*^ BMDMs (Fig 6b; Supplementary Fig 4). These results confirm that NLRP3 is responsible for sensing *S. aureus*, whereas AIM2 contributes minimally to this response.

**Fig 6:**
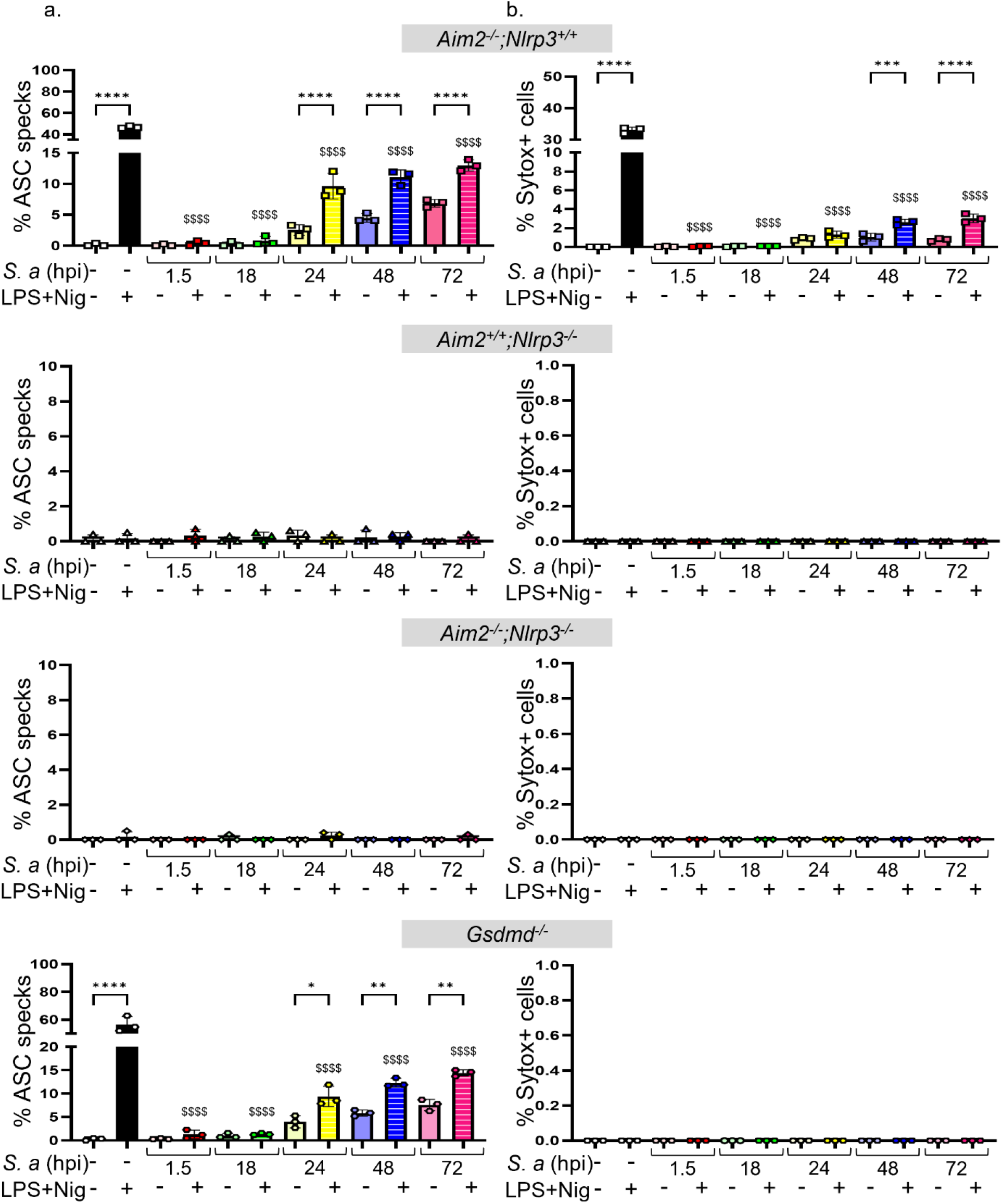
The formation of ASC specks and the uptake of Sytox green to a lesser extent, is induced by LPS and nigericin and depends on NLRP3. *Aim2*^*+/+*^*;Nlrp3*^*+/+*^, *Aim2*^*+/+*^*;Nlrp3*^*-/-*^, *Aim2*^*-/-*^*;Nlrp3*^*+/+*^, *Aim2*^*-/-*^*;Nlrp3*^*-/-*^, and *Gsdmd*^*-/-*^ BMDMs were left uninfected or infected with *S. aureus* for up to 72 h. a. ASC specks. b. Sytox Green. Data are means ± SD from experimental triplicates and represent at least two independent experiments. **p<0.01; ***p<0.001; ****p<0.0001 by one-way ANOVA with Bonferroni’s *post hoc* test. Comparison of LPS+Nig positive control with infection samples treated with LPS+Nig shown as $$$$ p<0.0001 by one-way ANOVA.

### Activation of the NLRP3 inflammasome is not restricted to BMDMs with *S. aureus* during late stages of infection

To further investigate the mechanisms underlying *S. aureus* regulation of the NLRP3 inflammasome pathway, we examined whether the presence of this bacterium inside the cell is necessary for ASC speck formation or Sytox Green uptake. Although the formation of ASC specks was marginal at the early phases of infection, up to 24 hpi, it was confined to *S. aureus*-engulfed BMDMs (Fig 7a). As the infection progressed, BMDMs without this bacterium also became ASC speck^+^ (Fig 7a). Intriguingly, Sytox Green uptake was predominantly restricted to BMDMs containing *S. aureus* (Fig 7b). These findings suggest that the internalization of *S. aureus* by BMDMs is essential for the initial activation of the inflammasome, however as the infection progresses, specks are also visible in cells cleared off bacteria too.

**Fig 7:**
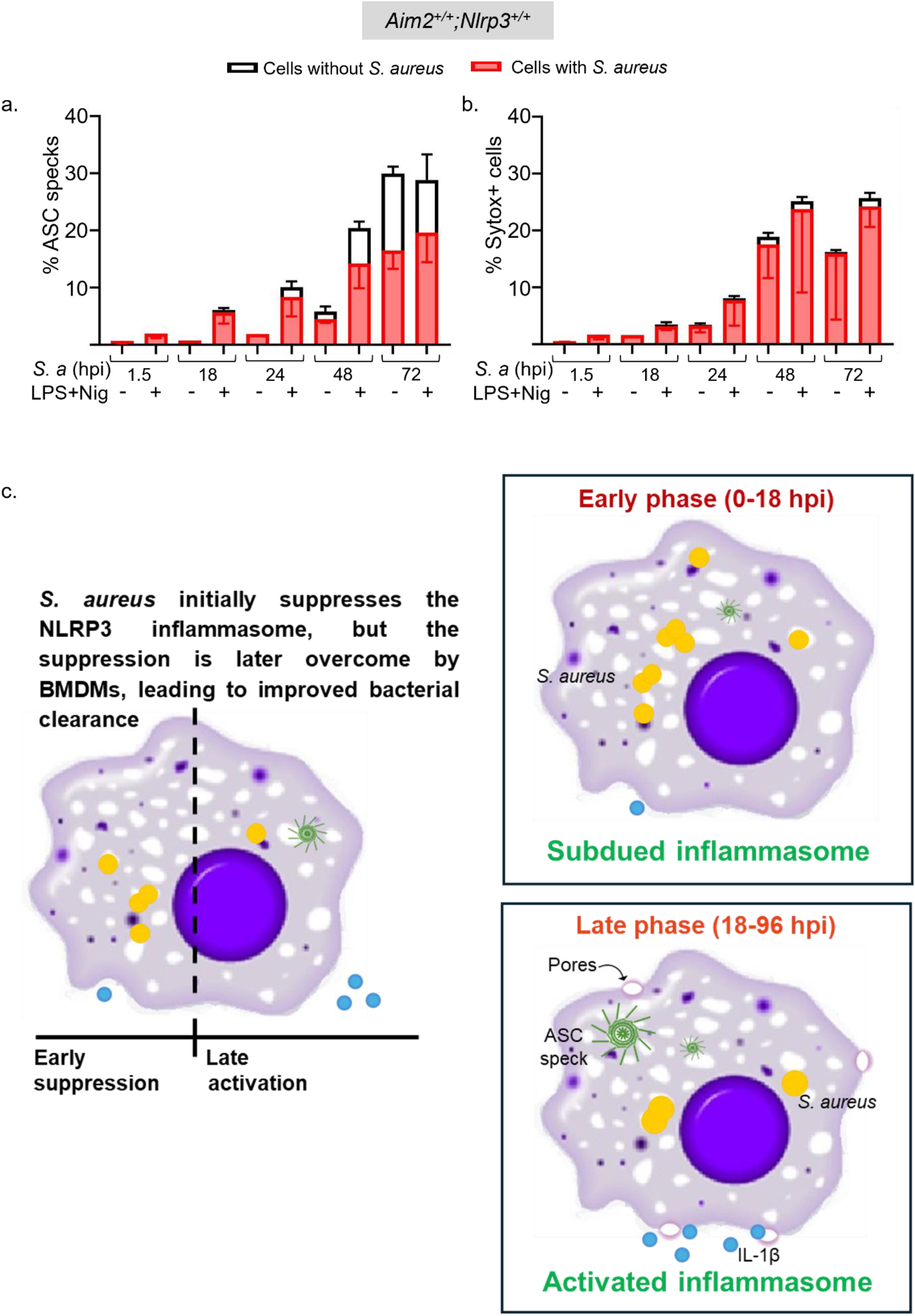
The NLRP3 inflammasome is suppressed during the early stages of *S. aureus* infection, but its activation is not restricted to BMDMs with internalized *S. aureus* during late stages of infection. BMDMs from *ASC-citrine* mice (to assess ASC speck formation) and WT mice (to measure Sytox Green uptake) were left uninfected or infected with *S. aureus* for up to 72 h were left untreated or treated sequentially with LPS for 3 h and nigericin for 45 min. a. ASC specks were quantified with respect to their colocalization with *S. aureus* signal within a BMDM. b. Sytox Green stained nuclei were quantified with respect to *S. aureus* signal within a BMDM. c. Graphical abstract. Data are means ± SD from experimental triplicates and represent at least two independent experiments.

## DISCUSSION

Previous studies indicated that NLRP3 and ASC were required for macrophages to activate caspase-1 and secrete IL-1β in response to signals induced by *S. aureus*^*14*^. Subsequent studies revealed that this inflammasome sensed *S. aureus* peptidoglycan, hemolysins, and lipoproteins^13,28,31^. In addition, caspase-1 was shown to promote phagosome-mediated killing of *S. aureus* through the acidification of these organelles^32^. In contrast, other data suggest that *S. aureus* virulence factors such as alpha toxin and peptidoglycan O-acetyltransferase A exploit both caspase-1 and caspase-11 to evade mitochondrial ROS-mediated killing^33–36^. Thus, conflicting findings have been reported regarding the role that inflammasomes play in determining the fate of *S. aureus* in macrophages. These discrepancies may stem from the *S. aureus* strains used and variations in experimental conditions, such as the duration of infection. In our time-course studies using a clinical *S. aureus* isolate, we found that the NLRP3 inflammasome can eventually overcome the evasive strategies through which *S. aureus* initially suppressed this pathway during the early stages of infection (Fig 7c). However, the specific mechanisms underlying this shift remain unclear. We also observed that ASC specks were initially detected only in infected BMDMs during the stages of infection, but they were later observed in both infected cells and *S. aureus*-free cells. It remains uncertain whether the infected cells eventually became pathogen-free or if ASC speck^+^ cells were bystanders responding to PAMPs or DAMPs. Regardless, these previously unreported biphasic responses underscore the dynamic nature of the interactions between *S. aureus* and host cells.

Although the NLRP3 inflammasome was progressively activated during *S. aureus* infection of BMDMs, as evidenced by ASC speck formation, caspase-1 activation, and IL-1β maturation, we found that GSDMD was not cleaved into its active p30 fragment during this process. It is possible that full-length GSDMD forms pores in the presence of *S. aureus*, as recently demonstrated in an experimental model unrelated to infection^37^. Alternatively, GSDMD might be post-translationally modified by the *S. aureus* strain used in this study. For instance, *Shigella flexneri* IpaH7.8 has been reported to ubiquitinate GSDMD in human epithelial cells, preventing its cleavage by caspase-1, as a strategy to maintain a niche for bacterial survival and growth^38^. The presence of several IL-1β species with higher molecular weight in *S. aureus*-infected BMDMs lends support to the hypothesis that this bacterium has the ability of modifying host’s effector molecules as reported by others^39^. Additionally, GSDMD-independent inflammasome mechanisms in host protection have been reported^40^. Contrasting studies also suggest that GSDMD-mediated pyroptosis plays a role in the pathogenesis of *S. aureus*-induced osteomyelitis^41^.

Our findings are primarily based on *in vitro* infections of BMDMs, which may not fully replicate the complexity of the *in vivo* bone marrow microenvironment or account for the contributions of other cell types involved in osteomyelitis. Despite these limitations, this study offers novel insights into the temporal regulation of NLRP3 inflammasome activity in BMDMs during *S. aureus* infection, uncovering a biphasic host response marked by early suppression followed by delayed activation.

## RESOURCE AVAILABILITY

## Lead contact

Requests for further information and resources should be directed to Gabriel Mbalaviele.

## Materials availability

This study did not generate new unique reagents

## Data and code availability

- Data reported in this paper will be shared by the lead contact upon request.
- This paper does not report any original code.
- Any additional information required to reanalyze the data reported in this paper is available from the lead contact upon request.

## ACKNOWLEDGMENTS

This work was supported by NIH/NIAID R01-AI161022, NIH/NIAMS R01 AR076758 and NIH/NIA R01 AG077732 (GM). We thank Crystal Idleburg and Samantha Coleman in Histology Core of Washington University Musculoskeletal Research Center (NIH P30 AR074992) for their assistance with confocal imaging. We would also like to thank Dr. Yousef Abu-Amer for granting permission to use the Flow Cytometer for our studies.

## AUTHOR CONTRIBUTIONS

Conceptualization, G.M., D.V., J.C., S.B.; methodology, S.B., K.K., C.W., G.M., D.V.; investigation, S.B.; analysis, S.B., G.S., G.M.; writing, S.B., G.M.; review, D.V., C.W., Y.L., K.K.; funding acquisition, G.M., D.V., J.C.; supervision, G.M.

## SUPPLEMENTAL INFORMATION

Figure Suppl 1; Figure Suppl 2; Figure Suppl 3-A; Figure Suppl 3-B; Figure Suppl 4-A; Figure Suppl 4-B

## STAR METHODS

### Study design

This study aimed to determine the response of BMDMs to *S. aureus* infection. To achieve this goal, we used genetically modified mouse strains and various biochemical and imaging approaches.

### Mice

To evaluate the role of inflammasomes in *S. aureus* infection, we purchased WT, R26-CAG-ASC-citrine (030744), *Aim2*^*-/-*^ (013144), and *Nlrp3*^*-/-*^ (021302) mice from The Jackson Laboratory (Sacramento, CA, USA). *Aim2*^*-/-*^ mice and *Nlrp3*^*-/-*^ mice were intercrossed to generate *Aim2*^*-/-*^ *;Nlrp3*^*-/-*^ mice. *Gsdmd* knockout (*Gsdmd*^−/−^) mice were kindly provided by Dr. V. M. Dixit (Genentech, South San Francisco, CA, United States of America). All mice were on the C57BL/6J background and mouse genotyping was performed by PCR as standardized before^42,43^. The Institutional Animal Care and Use Committee of Washington University School of Medicine in St. Louis approved all procedures.

### Bacterial strains and growth conditions

All experiments were conducted with derivatives of *S. aureus* USA300 clinical isolate Ti3^44^. Bacterial strains were grown in Trypticase soy broth (TSB) overnight at 37°C with shaking at 225 rpm, subcultured at a dilution of 1:100, grown to mid-exponential phase (optical density at 600 nm [OD_600_] of 1.0), and centrifuged at 3,000 rpm for 10 min. The pellets were washed and resuspended with PBS to the desired concentration.

### Cell cultures

BMDMs were obtained by culturing murine bone marrow cells in α-MEM culture media containing a 10% (v/v) conditioned media from the fibroblastic cell line CMG 14-12 as a source of macrophage colony-stimulating factor (M-CSF)^45^, for 4 to 5 days in a petri dish as previously described^46^. Briefly, nonadherent cells were removed by PBS washes and adherent BMDMs were detached with trypsin-EDTA and cultured in 10% CMG α-MEM culture media for various experiments. For all experiments, BMDMs were plated at 10^4^ cells per well on a 96-well plate or 10^6^ cells per well on a six-well plate.

### Serum optimization assay

WT BMDMs were cultured in media containing 0, 1, or 10% FBS and infected for 30 min at MOI of 1:100 at 37°C in 5% CO_2_, washed twice in PBS, and cultured in the same FBS-supplemented media containing the antibiotic, gentamicin (0.3 mg/ml) for 1 h to kill extracellular bacteria after which fresh media (α-MEM + M-CSF ± FBS without antibiotics) was added to the cells. Infection was quantified by CFU assay at 1.5 hpi and 18 hpi along the assessment of IL-1β secretion and cell death based on the release of lactate dehydrogenase (LDH). WT BMDMs cultured in different FBS concentrations were also subjected to known NLRP3 inflammasome activators, LPS and nigericin, as positive controls^47^.

### CFU assay

Infection of BMDMs by *S. aureus* was quantified by determining the number of colony-forming units (CFUs). Briefly, BMDMs from WT and different genotypes were seeded at 10^6^/well in six-well plates and cultured as described above. To determine the level of intracellular survival, the cells were infected for 30 min at an MOI of 1:100 at 37°C in 5% CO_2_, washed twice in PBS, and cultured in α-MEM media containing M-CSF and 0.3 mg/ml gentamicin for 1 h to kill extracellular bacteria. Cells were washed twice in PBS to remove the antibiotic and lysed in sterile, ice-cold ultrapure H_2_O at 1.5 hpi timepoint. For 18 hpi, culture media was replaced after PBS washes, and the infection was continued to 18 h before hypotonic lysis of the cells as described above. Lysates were 10-fold serially diluted, plated on TSB solidified with 1.5% agar (TSA), incubated overnight at 37°C, and CFU were enumerated. To ensure that extracellular *S. aureus* was effectively killed in all experiments, supernatants from antibiotic-treated cultures were plated on TSA and inspected for colonies after overnight incubation at 37°C.

### Western blot analysis

Cell extracts were prepared by lysing cells with radioimmunoprecipitation assay (RIPA) buffer (50 mM tris, 150 mM NaCl, 1 mM EDTA, 0.5% NaDOAc, 0.1% SDS, and 1.0% NP-40) and complete protease inhibitor cocktail. Protein concentrations were determined by the Bio-Rad Laboratories method, and equal amounts of proteins were subjected to SDS–polyacrylamide gel electrophoresis gels (12%) as previously described^43^. Proteins were transferred onto the PVDF membrane and incubated with antibodies against different protein components of the NLRP3 inflammasome pathway, NLRP3, GSDMD (1:1000; Abcam, MA), IL1β, α-tubulin and β-actin (1:2000; Santa Cruz Biotechnology, TX), overnight at 4°C followed by incubation for 1 h with respective secondary goat anti-mouse IRDye 800 (Thermo Fisher Scientific, MA) or goat anti-rabbit Alexa Fluor 680 (Thermo Fisher Scientific, MA). The results were visualized using the Odyssey Infrared Imaging System (LI-COR Biosciences, NE).

### LDH assay and IL-1β ELISA

Lytic cell death was assessed by the release of LDH in conditioned medium using the LDH Cytotoxicity Detection Kit (Takara, CA). IL-1β levels in conditioned media were measured by an ELISA kit (eBioscience, NY). For time-course infection analysis, culture supernatants were collected at each designated timepoint, after which fresh media was added to the cells and infection was continued until the next timepoint. Therefore, LDH and IL-1β levels measured at each timepoint represent the release relative to the preceding infection interval, rather than cumulative release.

### ASC specks assay

*Aim2*^*+/+*^*;Nlrp3*^*+/+*^, *Aim2*^*+/+*^*;Nlrp3*^*-/-*^, *Aim2*^*-/-*^*;Nlrp3*^*+/+*^, *Aim2*^*-/-*^*;Nlrp3*^*-/-*^, and *Gsdmd*^*-/-*^ BMDMs were seeded overnight and infected with *S. aureus* at 100 MOI and/or stimulated for inflammasome activation with LPS for 3 h before adding 15 μM nigericin for 45 min as previously described^47^. After the appropriate treatments and infection timepoints, cell culture supernatants were collected, and the cells were washed with PBS and fixed with 4% paraformaldehyde buffer for 10 min at room temperature, followed by permeabilization and staining for ASC using anti-mouse ASC antibody (EMD Millipore) diluted in buffer containing 0.2% triton and 1% BSA in PBS. After overnight incubation at 4°C, cells were stained with a secondary antibody (Alexa Fluor 594; Life Technologies) for 30min and counterstained with Fluoro-gel II containing 4′,6-diamidino-2-phenylindole (Fluoro-Gel, Fisher Scientific Intl. Inc., PA). The formation of ASC specks, indicating inflammasome assembly, was detected using a Leica inverted microscope with a TCS SPE II confocal module and processed using LAS X software (Leica Microsystems Inc., IL). Quantification of ASC specks was conducted using ImageJ.

### Sytox Green uptake assay

BMDMs were seeded in tissue culture plates and either primed using LPS or given *S. aureus* infection for the indicate timepoints. The cells were incubated with Hoechst 33342 (1 µg/mL) for 15 min, the media was changed, and treated with nigericin for 45 min. Following stimulation, Sytox Green (Invitrogen) diluted in PBS was added to the wells to reach a final concentration of 10 nM. The cells were incubated for 10-15 min to allow the dye to penetrate cells. Fluorescence was measured in live cells using a Leica inverted microscope with a TCS SPE II confocal module and processed using LAS X software (Leica Microsystems Inc., IL). The percentage of Sytox Green-positive cells was analyzed and quantified using ImageJ.

### Flow cytometry

BMDMs were seeded in 6-well tissue culture plates and challenged with GFP-expressing Ti3-*S. aureus* at an MOI of 1:100 for 30 min. Extracellular bacteria were killed by the addition of gentamicin to the cultures for 1 h. Cells were washed twice in PBS to remove the antibiotic, the media was replenished and harvested at the indicated timepoints. Cells were detached with trypsin-EDTA, washed twice in PBS to remove nonadherent bacteria as previously described^48^, and analyzed by flow cytometry (percent fluorescein isothiocyanate [FITC]-positive cells and mean fluorescence intensity of FITC-positive population). Flow cytometry acquisition was performed using the Attune Flow Cytometer system, followed by analysis with FlowJo software (Tree Star, Ashland, Oregon).

### FLICA assay

Active caspase-1 was labeled using the FAM-FLICA Caspase-1 (YVAD) Assay Kit (Immunochemistry Technologies, MN). BMDMs were seeded as described above and stimulated with *S. aureus* in the absence or presence of LPS and nigericin, then treated with 10 μl of 30X FLICA solution. The cells were stained with Hoechst 33342 stain (0.5% w/v) and viewed using a Leica inverted microscope with excitation 490 nm and emission >520 nm for Green fluorescence and excitation 365 nm and emission 480 nm for the Hoechst dye. The percentage of cells staining Green (indicating active caspase-1) were quantified using ImageJ software.

## Statistical analysis

Statistical analysis was performed using the Student’s *t* test or one-way analysis of variance (ANOVA) with Bonferroni’s multiple comparisons test using the GraphPad Prism Software. Values are expressed as means ± SEM or means ± SD, as indicated. *p<0.05 was considered statistically significant.

