## Supplementary figures for "*Staphylococcus aureus* inhibits the NLRP3 inflammasome in macrophages during the early phases of intracellular infection, but not the late phases"

Bhagat, et. al.

**Supplementary Figure Legends:**

**Suppl 1: Flow cytometric analysis of *S. aureus*-infected BMDMs.** *Aim2+/+;Nlrp3+/+*, *Aim2+/+;Nlrp3-/-, Aim2-/-;Nlrp3+/+,* and *Aim2-/-;Nlrp3-/-* BMDMs were infected with GFP-tagged *S. aureus.* a. Flow cytometry analysis. b. quantitative tabulation of the frequencies of GFP+ cells with respect to total live cells. Data are average values from three biological replicates.

**Suppl 2: *S. aureus* suppresses the activation of caspase-1 during the early phases of infection of BMDMs, but to a lesser extent during the late phases -** BMDMs were infected with *S. aureus* for up to 48 h. Cells left untreated or treated sequentially with LPS for 3 h and nigericin for 45 min then incubated with 10 µl of 30X FLICA solution followed by the staining with Hoechst 33342. a. FLICA b. Quantitative data. *n* = 3 technical replicates, representative of 3 biological replicates.

**Suppl 3: The formation of ASC specks induced by LPS and nigericin depends on NLRP3. S3-A:** *Aim2+/+;Nlrp3-/-, Aim2-/-;Nlrp3+/+,* **S3-B:** *Aim2-/-;Nlrp3-/-,* and *Gsdmd-/-* BMDMs were left uninfected or infected with *S. aureus* for up to 72 h. Cells were left untreated or treated sequentially with LPS for 3 h and nigericin for 45 min. ASC specks were visualized under fluorescence microscopy.

**Suppl 4: The uptake of Sytox green induced by LPS and nigericin depends on NLRP3. S4-A:** *Aim2+/+;Nlrp3-/-, Aim2-/-;Nlrp3+/+,* **S4-B:** *Aim2-/-;Nlrp3-/-,* and *Gsdmd-/-* BMDMs were left uninfected or infected with *S. aureus* for up to 72 h. Cells were left untreated or treated sequentially with LPS for 3 h and nigericin for 45 min. Sytox green was added to the cultures 10 min prior to imaging by confocal microscopy. Data is representative of at least two independent experiments.

### Suppl 1

a.

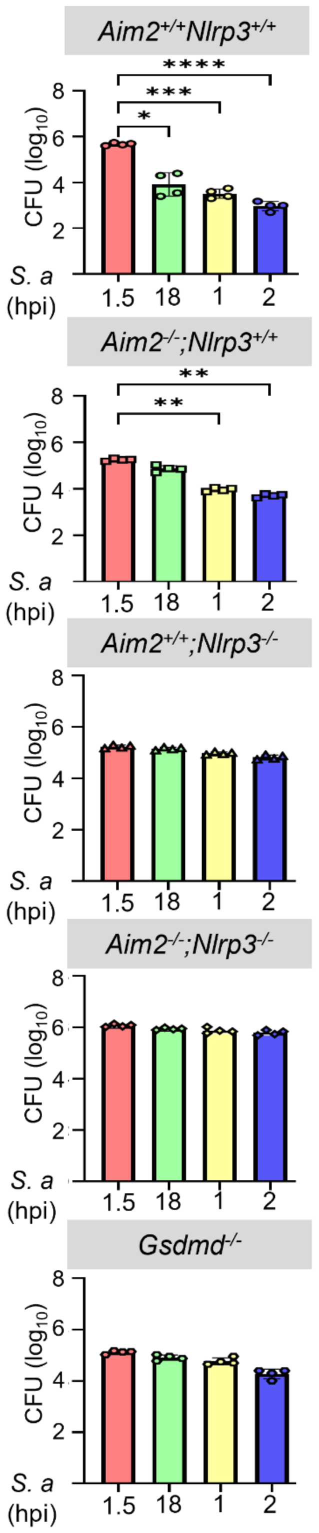


b.

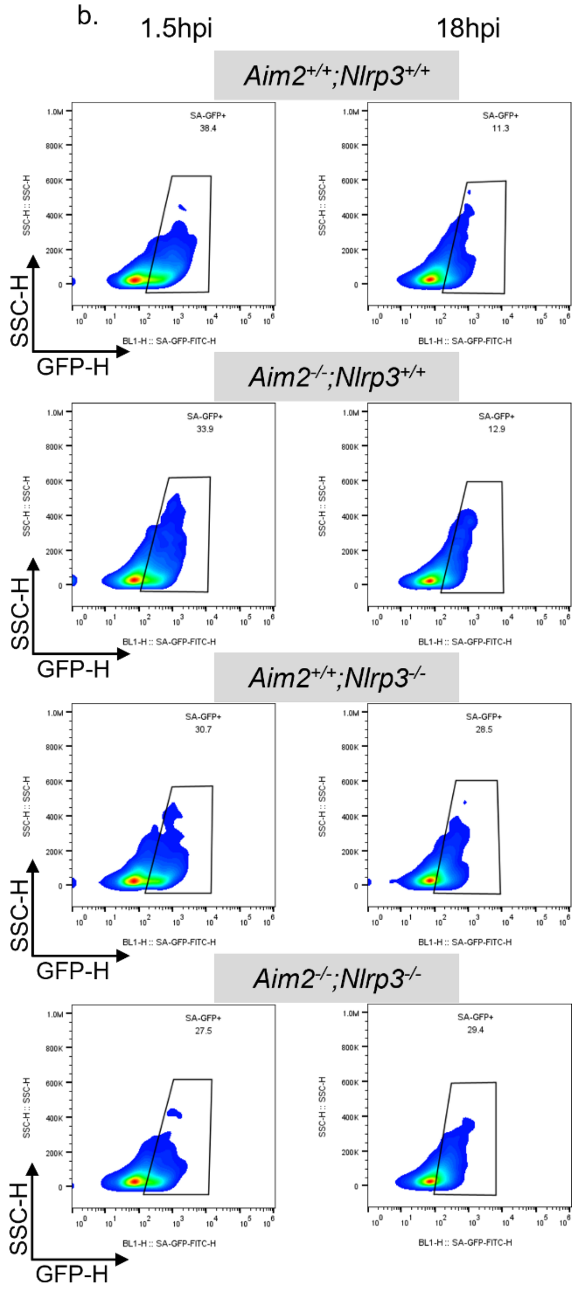


c.

| **Genotype** | ***S. a* infection** | |
|---|---|---|
| | **Freq of GFP+ cells** | |
| | 1.5hpi | 18hpi |
| *Aim2+/+;Nlrp3+/+* | 37.4 | 13.5 |
| *Aim2-/-;Nlrp3+/+* | 39.9 | 13.2 |
| *Aim2+/+;Nlrp3-/-* | 30.4 | 29.2 |
| *Aim2-/-;Nlrp3-/-* | 30.8 | 28.0 |

### Suppl 2

*Aim2^+/+;Nlrp3^+/+*

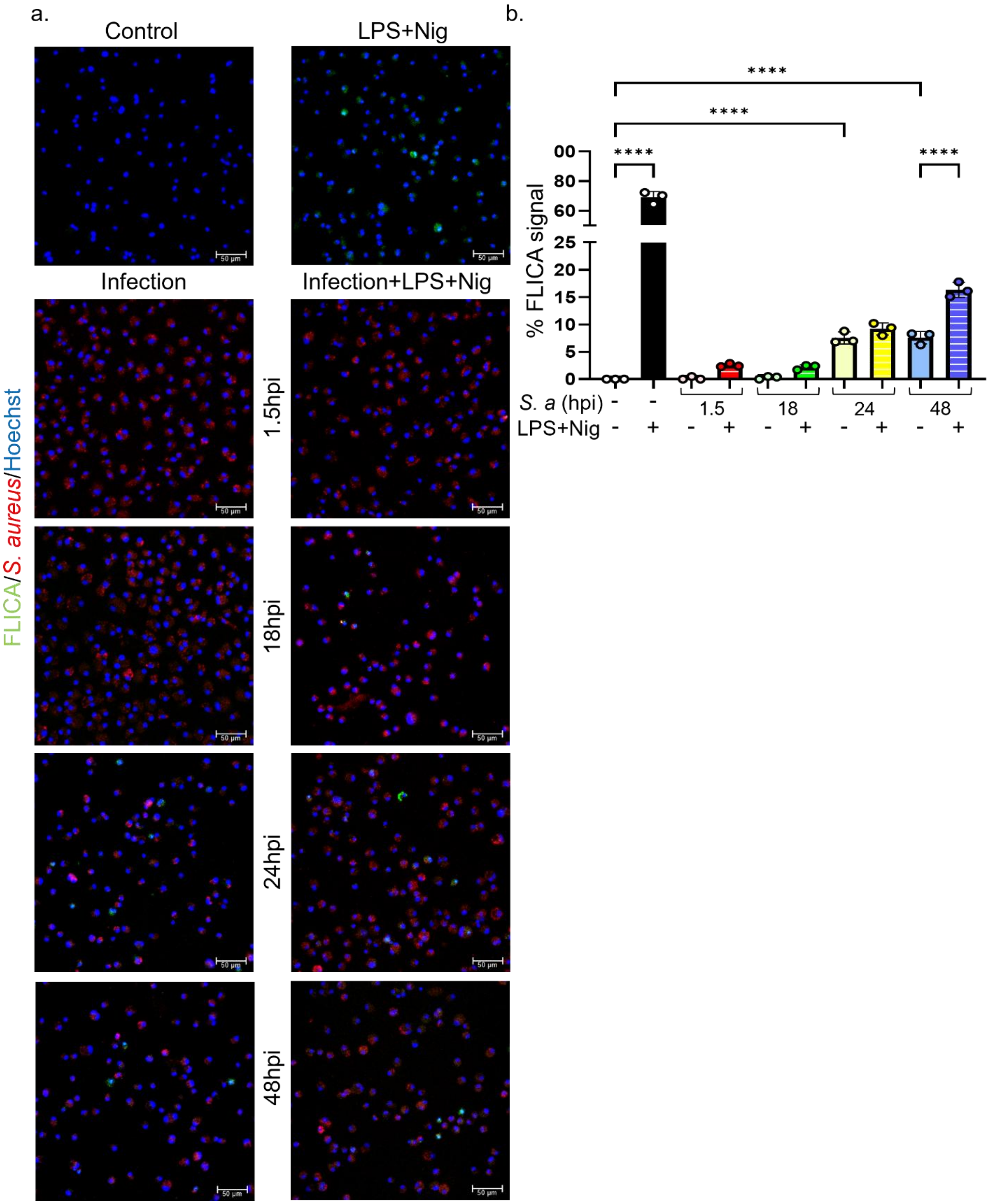

### Suppl 3

S3-A.

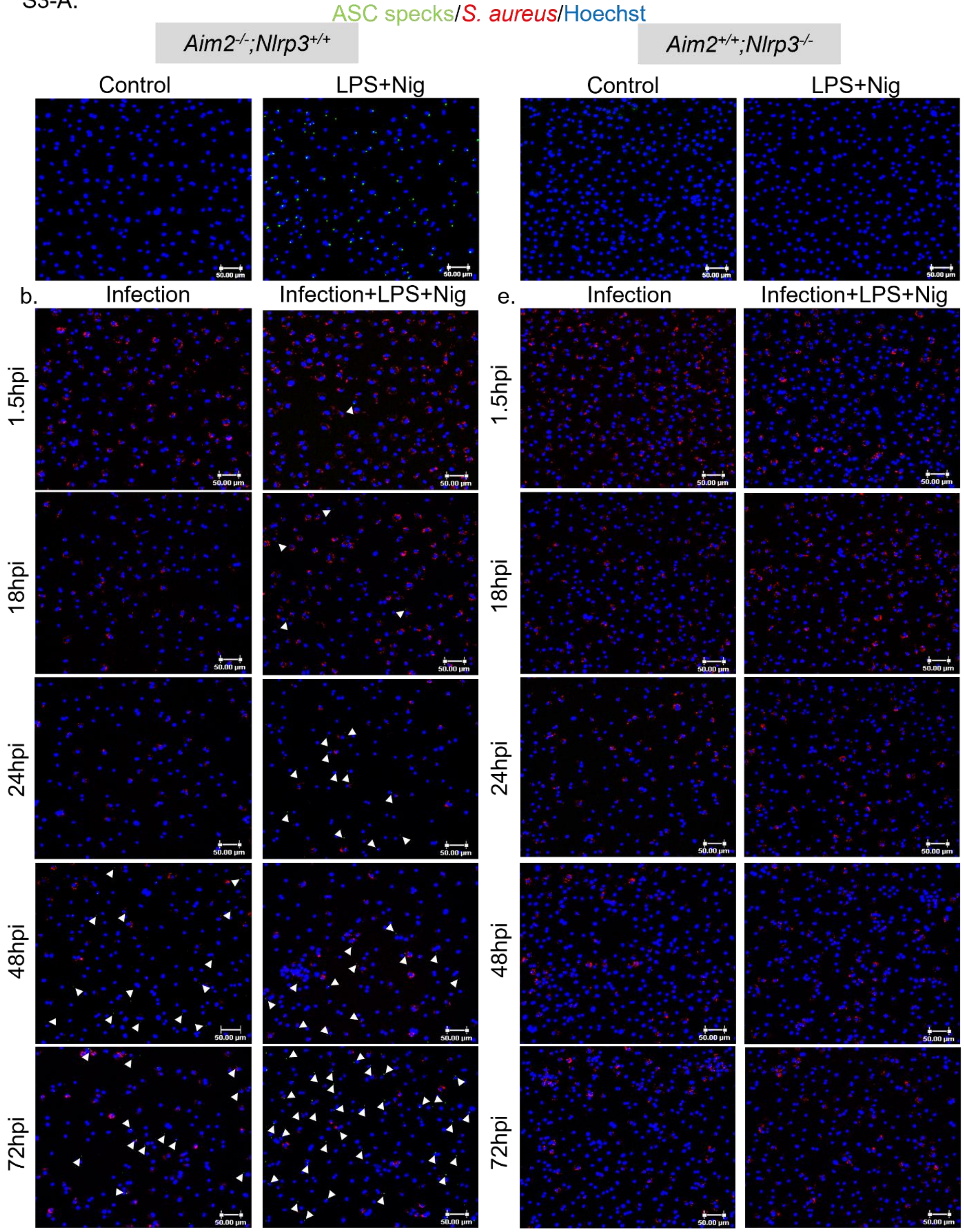

ASC specks/*S. aureus*/Hoechst
*Aim2*-/-;*Nlrp3*+/+
*Aim2*+/+;*Nlrp3*-/-
Control
LPS+Nig
Control
LPS+Nig
b.
Infection
Infection+LPS+Nig
e.
Infection
Infection+LPS+Nig
1.5hpi
18hpi
24hpi
48hpi
72hpi
1.5hpi
18hpi
24hpi
48hpi
72hpi
50.00 μm

### Suppl 3

S3-B.

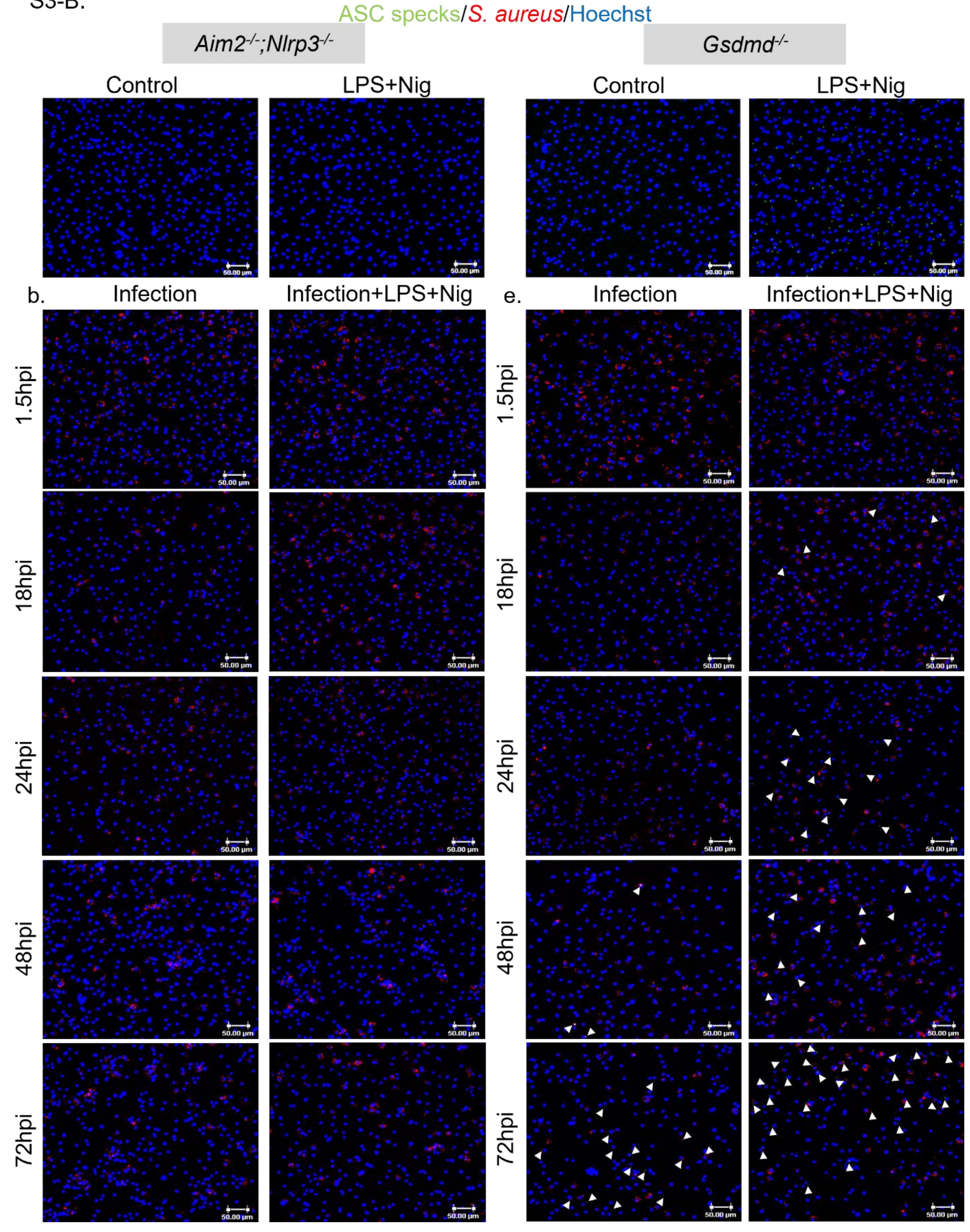
ASC specks/S. aureus/Hoechst
Aim2-/-;Nlrp3-/-
Gsdmd-/-
Control
LPS+Nig
Control
LPS+Nig
b.
Infection
Infection+LPS+Nig
e.
Infection
Infection+LPS+Nig
1.5hpi
18hpi
24hpi
48hpi
72hpi
50.00 µm

### Suppl 4

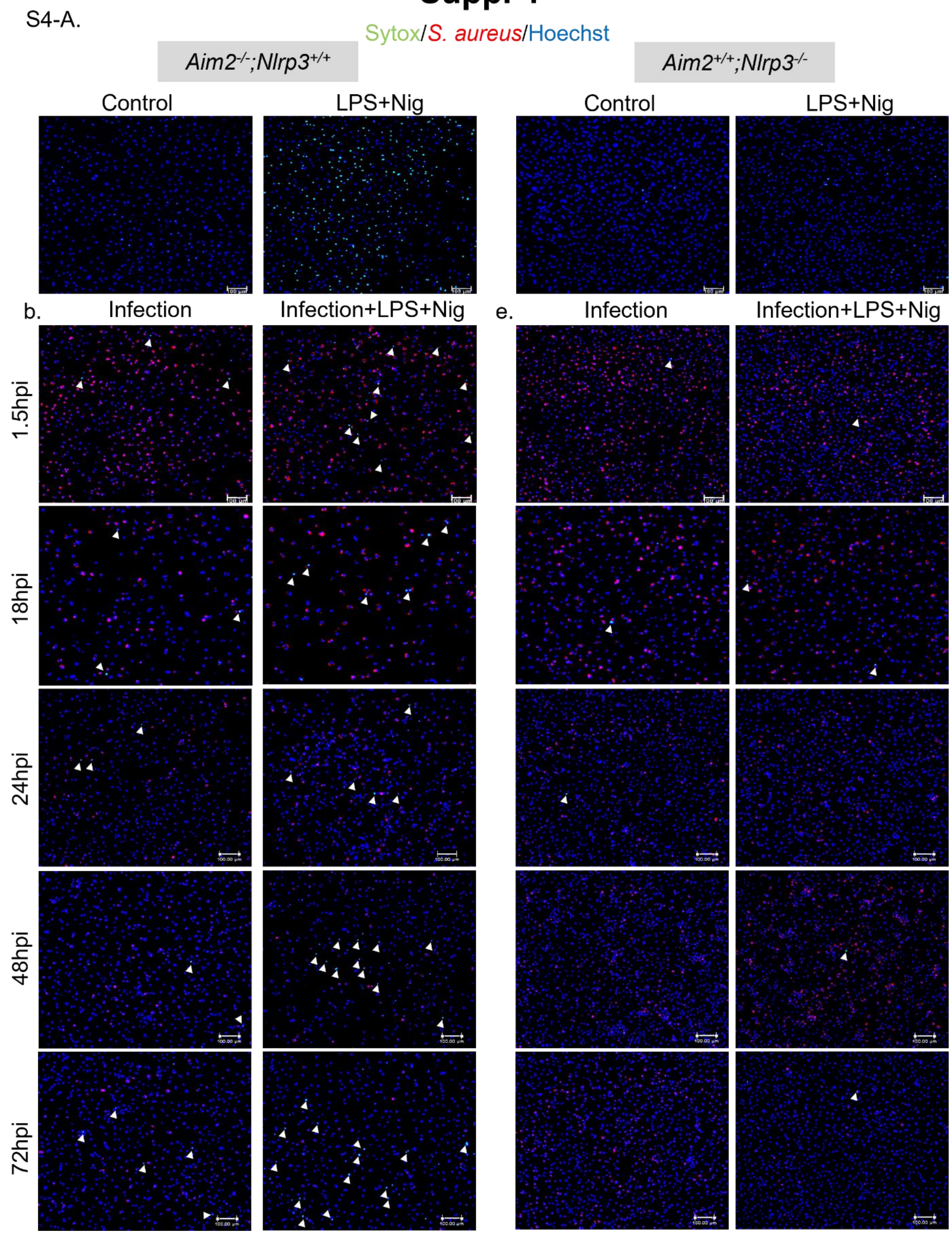

S4-A.
Sytox/S. aureus/Hoechst
Aim2-/-;Nlrp3+/+
Aim2+/+;Nlrp3-/-
Control
LPS+Nig
Control
LPS+Nig
b.
Infection
Infection+LPS+Nig
e.
Infection
Infection+LPS+Nig
1.5hpi
18hpi
24hpi
48hpi
72hpi

### Suppl 4

S4-B.

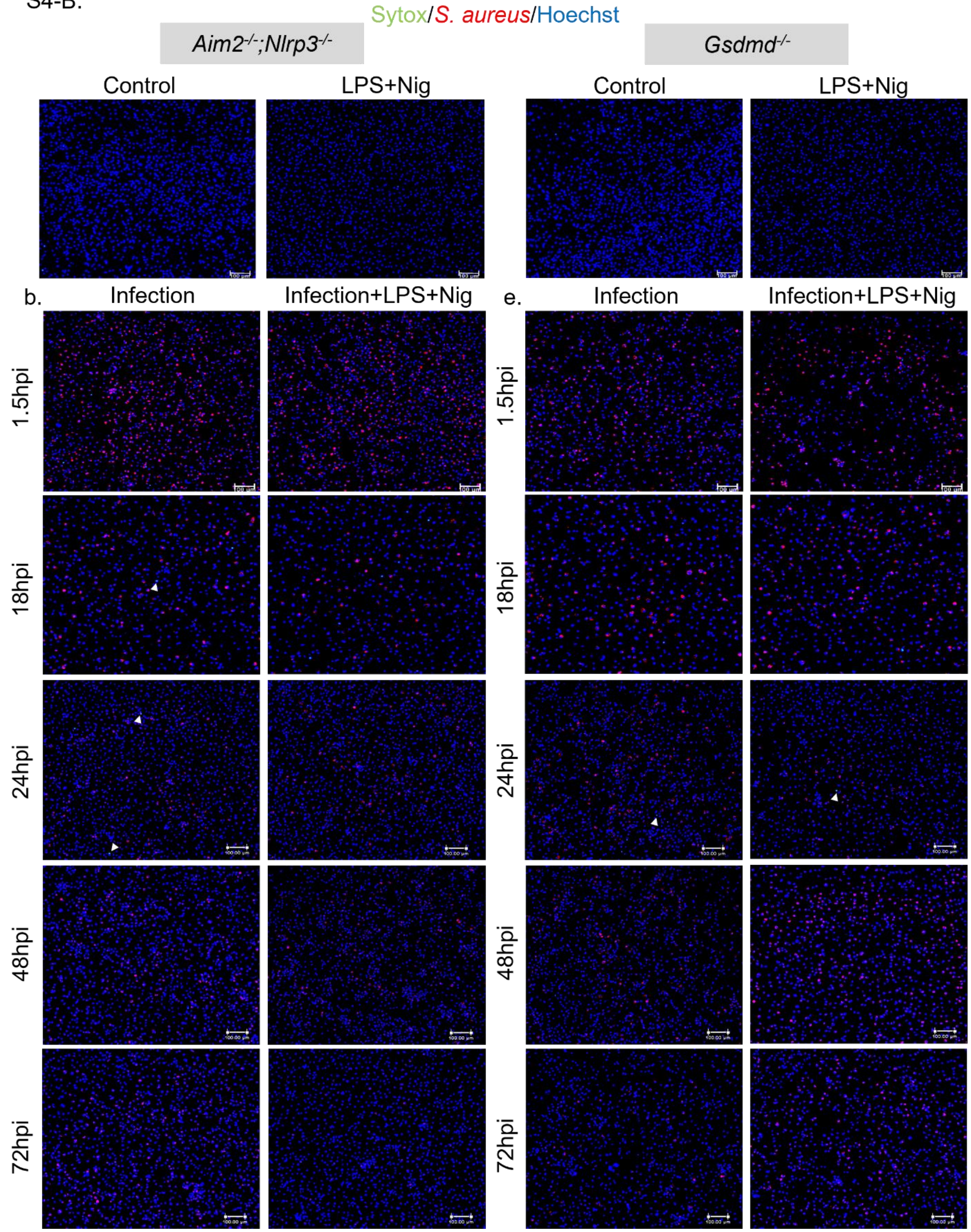

Sytox/S. aureus/Hoechst
Aim2-/-;Nlrp3-/-
Gsdmd-/-
Control
LPS+Nig
Control
LPS+Nig
b.
Infection
Infection+LPS+Nig
e.
Infection
Infection+LPS+Nig
1.5hpi
18hpi
24hpi
48hpi
72hpi
1.5hpi
18hpi
24hpi
48hpi
72hpi
